# Phylogeny-informed transfer learning with protein language models for epitope prediction

**DOI:** 10.1101/2025.04.17.649425

**Authors:** Lindeberg Pessoa Leite, Teófilo Emidio de Campos, Francisco Pereira Lobo, Felipe Campelo

## Abstract

Generalist linear B-cell epitope predictors are typically trained on large, heterogeneous datasets, which can lead to biased representations and degraded performance for under-represented or emerging pathogens. We present a transfer learning framework leveraging the ESM family of protein language models (PLMs) as sequence feature embedders, adapting them to specific evolutionary contexts through phylogeny-informed fine-tuning. By coupling pretrained PLM representations with hierarchical, taxon-aware adaptation, our approach enables efficient knowledge transfer from sets of related pathogens to data-scarce targets while preserving lineage-specific signals. This strategy consistently improves predictive performance over state-of-the-art epitope prediction methods across a diverse set of targets. We further demonstrate that these gains arise from the targeted adaptation and downstream models guided by evolutionary relatedness, indicating the value of this structured transfer learning for epitope prediction and highlighting its potential usability for further predictive applications with evolutionarily-structured data.

## 1 Introduction

Deep learning methods are increasingly used for modelling protein sequences (Collatz et al., 2020; Bahai et al., 2021; Rives et al., 2021; Croce et al., 2024), with transfer learning approaches that leverage large pretrained protein language models (PLMs) emerging as particularly effective (Fenoy et al., 2022; Vu et al., 2024; Liu et al., 2024; Schmirler et al., 2024). PLM-derived latent representations capture rich, context-dependent biological informationp (Rives et al., 2021; Elnaggar et al., 2021; Chowdhury et al., 2022; Lin et al., 2023) and have enabled substantial progress across downstream tasks (Schmirler et al., 2024). One such task is the computational prediction of linear B-cell epitopes (LBCEs) (Clifford et al., 2022; Gabellieri et al., 2026), an important step in a broad range of medical and immunological applications including the development of vaccines (Hamley, 2022), therapeutic antibodies (Sun et al., 2024), and immunodiagnostics (Mucci et al., 2017).

Most existing LBCE prediction approaches, including recent methods leveraging PLMs (Clifford et al., 2022; Høie et al., 2024), are trained using labelled peptide sequences from a phylogenetically diverse range of organisms. As we have previously discussed (Ashford et al., 2021; Campelo and Lobo, 2024), the main goal of epitope prediction methods developed using heterogeneous datasets is to provide general-purpose models that can be readily used without referencing the source organism of the proteins or peptides queried, an approach that can obscure lineage-specific signals and reduce performance for neglected or under-studied pathogens.

Building on evidence that taxon-aware modelling improves target-specific accuracy (Ashford et al., 2021; Lim et al., 2021; Yin et al., 2022; da Silva et al., 2023; Liu et al., 2023; Campelo and Lobo, 2024; Campelo et al., 2024), we develop a phylogeny-informed transfer learning framework that fine-tunes general-purpose PLM embedders using data from evolutionarily related organisms, enabling structured adaptation of learned representations to the target lineage. This strategy yields substantial and consistent performance gains across diverse pathogens, over both internal baselines and external, state-of-the-art predictors, demonstrating that the incorporation of evolutionary relationships provides a principled and effective mechanism for adapting PLMs to epitope prediction and related genomic modelling tasks.

## 2 Results

The central objective of this work is to investigate the effect of phylogeny-informed transfer learning (PITL) on the performance of downstream predictive tasks, using the LBCE prediction problem as a case study of particular relevance. To this end, we developed a modular phylogeny-informed modelling approach for the generation of taxon-specific linear B-cell epitope predictors (Ashford et al., 2021; Campelo et al., 2024). The data sources and modelling steps are outlined in Figure 1. The performance of this approach was assessed using a phylogenetically diverse set of nineteen target taxa of interest including viruses, bacteria and eukaryotic pathogens. Details regarding these datasets, including data retrieval, quality control, data consolidation, and the total volume of available data under each dataset are provided in the Supplementary File (Section “Data”) and Supplementary Tables 1 – 2.

**Figure 1.**
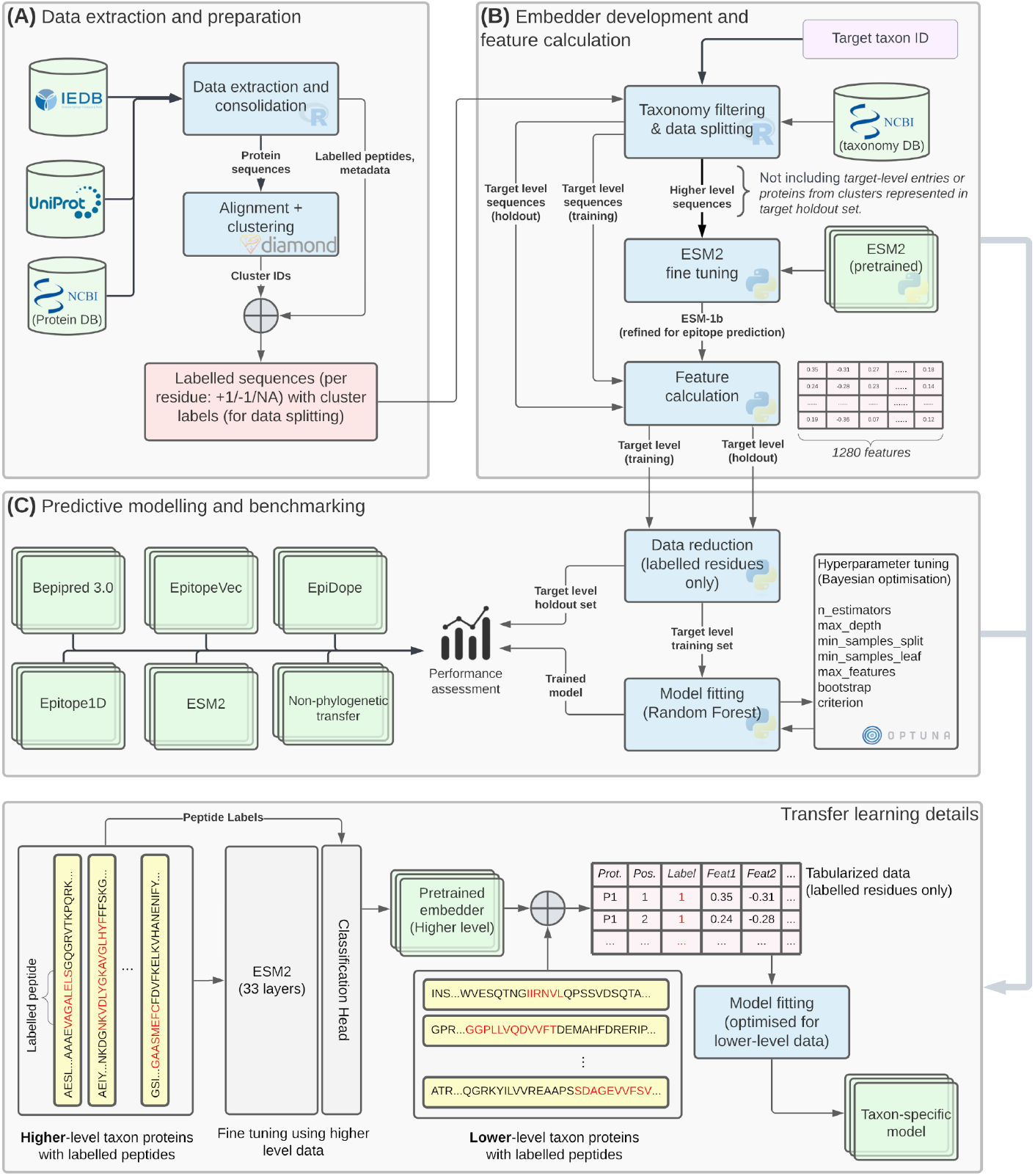
Overview of the framework for building taxon-specific LBCE predictors with phylogeny-informed transfer learning. **(A)** Retrieval of labelled peptides and their source protein sequences. Protein sequences are clustered based on normalised alignment scores, using single-linkage hierarchical clustering with a local similarity threshold of 30%. The resulting clusters are treated as allocation units when splitting data for model development/training and testing to prevent data leakage due to sequence similarity. **(B)** Filtering of the consolidated data according to the target taxon of interest. Entries associated with a higher taxonomic level (e.g., kingdom or phylum; excluding all entries from the target taxon itself) are used to fine tune the ESM feature embedder, which is then applied to extract features for entries belonging to the target taxon. **(C)** Model fitting. Data related to the target taxon (e.g., species or genus) is used to fit and optimise a classifier specifically for the target level. The effects of the transfer learning strategy are evaluated by benchmarking the final models on holdout sets (completely unseen during embedder fine tuning or model development). Their performance is compared against models trained using the baseline ESM features (without fine-tuning), and using an ESM variant tuned using epitope data from pathogens without any close phylogenetic link to the target taxon; and against external benchmarks.

We created two sets of internal baselines to investigate the effect of the proposed approach on predictive performance. The first employs the general-purpose PLM without any fine tuning (no Transfer Learning, *NTL*). This baseline allows the estimation of the effect of embedder fine tuning, by contrasting their predictive performance against models developed based on PITL. The second internal baseline uses *phylogeny-agnostic transfer learning* (*PATL*), in which the PLM is fine tuned using data of pathogens from domains distinct from the target taxon, i.e., with negligible phylogenetic links (Supplementary Figure 1). Since this second baseline differs from PITL only by not considering phylogenetic links in the fine-tuning step, it allows the estimation of the specific effect of using phylogeny as a criterion when selecting data for the fine-tuning step.

In addition to the internal baselines listed above, four external comparison baselines were used: BepiPred 3.0 (Clifford et al., 2022), Epidope (Collatz et al., 2020) and EpitopeVec (Bahai et al., 2021), which are deep learning-based generalist approaches using sophisticated feature spaces for LBCE prediction; and epitope1D (da Silva et al., 2023), a recent approach which provides taxon-specific models, commonly stratified at the phylum or kingdom level. These external baselines were selected as representative of the current state-of-the-art in LBCE prediction which provide online interfaces or easily reproducible source code.

### 2.1 Phylogeny-informed transfer learning improves LBCE prediction

The results indicated significant differences in the performance of the methods (ANOVA *p* = 0.0035), but not between ESM versions (*p* = 0.3183). In the pairwise comparisons we observed statistically significant AUC gains of PITL-based models in relation to both internal baselines. When compared against the NTL baseline, PITL-based models display significant gains (Dunnett *p* = 0.004; *CI*_95%_ = 0.032 *±* 0.023; Cohen’s *d* = 0.72; Figure 2(a)). Relevant effect sizes were also observed for other performance indicators (Supplementary Table 3), such as the Matthews Correlation Coefficient (MCC; *p* = 0.0016; *CI*_95%_ = 0.095 *±* 0.062; *d* = 0.79), a robust, class imbalance-resistant performance indicator (Chicco and Jurman, 2020). These results indicate that the fine-tuning step by itself results in performance gains for the downstream predictive task, well balanced across positive and negative predictions.

**Figure 2.**
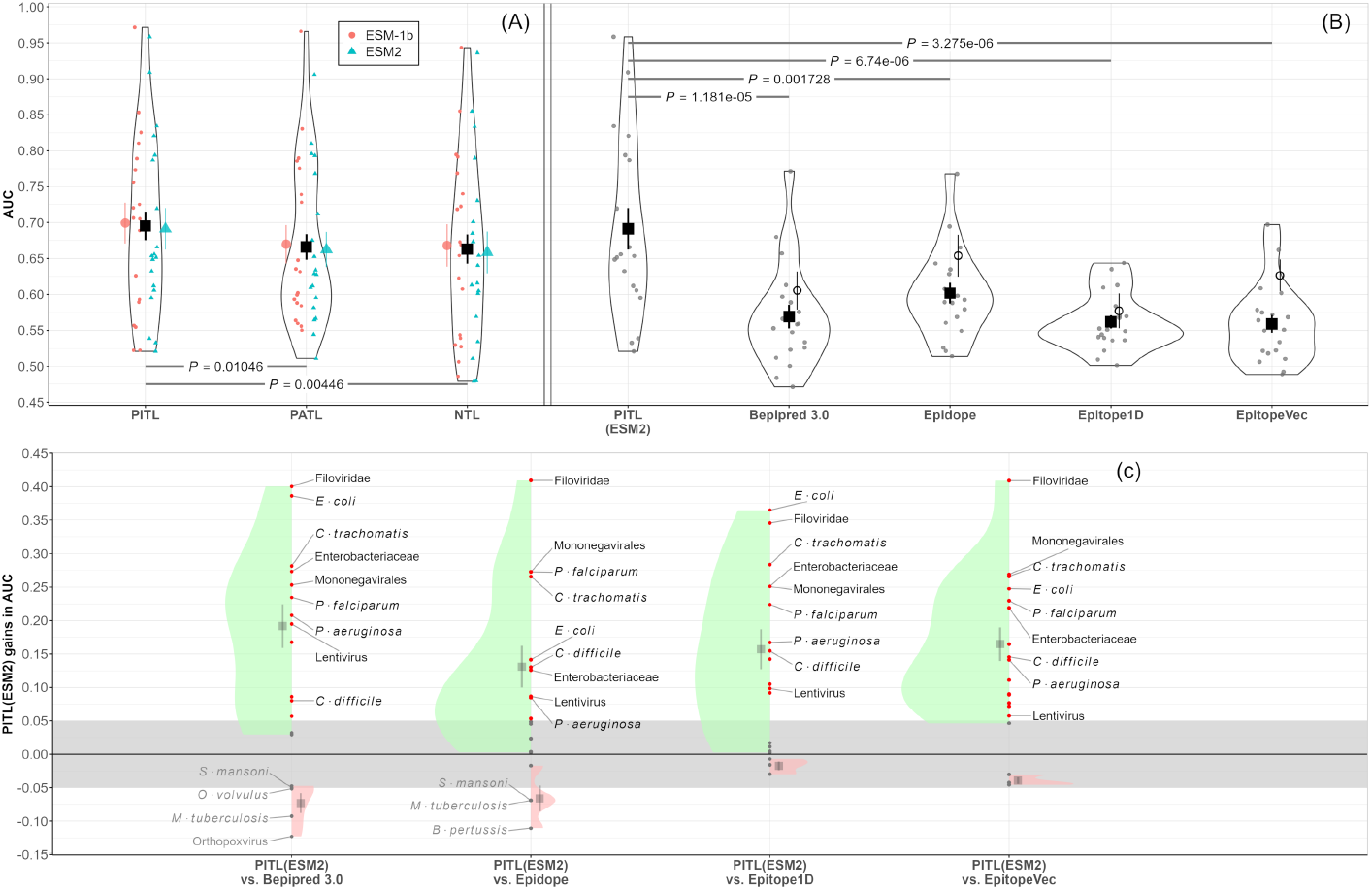
omparison of PITL-based models against internal and external baselines. **(A)** PITL-based models provide significant AUC gains when compared to similar models built using PATL- or NTL-based embeddings. Small points represent AUC values observed for individual datasets. Larger points with standard error bars represent estimates of mean performance for method/embedder version combinations. The p-values correspond to post-ANOVA pairwise comparisons between PITL and the internal baselines, after accounting for the effects of embedder version and dataset. **(B)** Comparison of the PITL(ESM2) against external baselines. Small points indicate performance estimates on individual datasets, larger square markers represent mean performances obtained considering the *No-leakage holdout*, with associated standard errors. For the external baselines, the mean and standard errors calculated using the *Internal holdout* are also displayed as hollow circles, for reference only. **(C)**: Distribution of per-dataset performance gains of PITL(ESM2)-based predictors in relation to the external baselines, illustrating the difference between observed performance gains (green) and losses (red). Differences of ±0.05 in AUC, shown as a grey band, were deemed to be of practical equivalence. Mean gains and mean losses are displayed as grey squares with associated standard errors. PITL(ESM2)-based models that presented gains greater than *±*0.05 against all baselines are explicitly labelled, as well as all cases of performance loss outside the region of practical equivalence. See section “Methods - Performance assessment and comparison” for details on the two distinct definitions of hold-out set used.

One of the core hypotheses of this work is that data from phylogenetically closer pathogens carries more relevant information for training taxon-specific predictors than data from more distantly related organisms. This was tested by contrasting the performance of PITL-based models with their PATL-based counterparts (see section “Methods - Performance assessment and comparison”; and Supplementary Figure 1). The results obtained in this comparison indicate clear gains in performance resulting from the incorporation of a phylogeny-informed strategy for the embedder fine-tuning step (Figure 2(b)). We observed a statistically significant improvement in AUC (Dunnett *p* = 0.0105; *CI*_95%_ = 0.029 *±* 0.023; *d* = 0.65; Figure 2(a)) as well as in other performance indicators such as the MCC (*p* = 0.0004; *CI*_95%_ = 0.107 *±* 0.062; *d* = 0.89). Taken in combination with the observed improvement over the NTL-based baselines, these results provide compelling evidence that the superior performance of PITL-based models can be specifically attributed to the use of data from pathogens that are phylogenetically close to the target taxon for fine-tuning the ESM embedder, rather than it being the effect of simply refining the PLM for the general epitope prediction task.

### 2.2 Models based on PITL(ESM2) show better predictive performance than state-of-the-art approaches

We selected PITL(ESM2)-based models for further comparisons with external baselines, and compared the results obtained by these models against four state-of-the-art LBCE predictors. Three of those are deep learning-based, generalist LCBE predictors: BepiPred 3.0 (Clifford et al., 2022), which employs NTL(ESM2) embeddings augmented by surface accessibility and protein length features; Epidope (Collatz et al., 2020), which utilises a deep neural network sequence embedder known as ElMo (Heinzinger et al., 2019); and EpitopeVec (Bahai et al., 2021), which is based on a combination of amino acid features, modified antigenicity scales and neural network-based protein embeddings. The fourth external comparison baseline, Epitope1D (da Silva et al., 2023), implements a rough taxon-specific modelling approach, with models provided for major phyla or kingdoms of bacterial, viral, and eukaryotic pathogens. Epitope1D’s feature space is based on a bespoke graph-based attributes extracted from a set of physicochemical and statistical amino acid descriptors.

The main purpose of these comparisons is to establish whether the specific combination of components used in the PITL-based modelling pipeline is sufficient to produce predictors with an expected performance comparable or superior to the best current predictive methods. As shown in Figure 2(B), we found statistically significant differences in mean AUC between PITL(ESM2) and all four external baselines, with effect sizes noticeably greater than the ones observed in the comparisons against the internal baselines. The observed performance gains of PITL(ESM2)-based models were substantial in relation to the three generalist baselines: Epidope (Dunnett *p* = 0.0017; *CI*_95%_ = 0.130 *±* 0.061); *d* = 1.19), BepiPred 3 (*p* = 1.10*×*10^−5^; *CI*_95%_ = 0.090*±*0.061; *d* = 1.62) and EpitopeVec (*p* = 1.86*×*10^−6^; *CI*_95%_ = 0.120*±*0.061; *d* = 1.76). Substantive AUC gains were also found in the comparison against Epitope1D, which also uses taxon-specific models albeit at a coarser granularity (*p* = 3.70*×*10^−6^; *CI*_95%_ = 0.132 *±* 0.061; *d* = 1.72). Supplementary Table 4 provides the full description of the comparisons across multiple performance indicators.

Figure 2(C) displays the distribution of the magnitude of paired differences of AUC for each dataset, stratified in terms of performance gains/losses of PITL(ESM2)-based models in relation to the external baselines. Consistently with the results above, we observe performance gains in the majority of the 19 comparisons, with a few instances displaying performance degradation against BepiPred 3 and Epidope (four and three cases, respectively). The average magnitude of gains are also substantially greater than those of performance decreases. For twelve of the nineteen datasets used in this experiment, PITL(ESM2)-based models exhibited positive AUC gains against every external baseline; in nine of those, the AUC gains were all greater than our (arbitrarily defined) minimally relevant effect size of +0.05. These include models across the three domains explored: viral, bacterial, and eukaryotic pathogens. Five of these models exhibiting relevant AUC gains correspond to specific pathogens; the other four were derived for higher level target taxa, such as genus or family.

### 2.3 Use of PITL results in high-performance LBCE predictors across taxonomically diverse pathogens

Some of the predictors instantiated in this study present particularly high predictive performance, making them potential models of choice for researchers investigating certain pathogens. The model for the Filoviridae family, which contains highly virulent pathogens such as the Ebola and Marburg viruses (Kuhn et al., 2019), is the one where observed performance differences are most prominent. As shown in Figure 2(C), this model resulted in absolute gains in AUC greater than 0.4 in relation to BepiPred 3, EpitopeVec and Epidope, and greater than 0.35 in comparison to Epitope1D. Supplementary Table 5 reports the full performance values observed for this model, which displays an out-of-sample AUC of 0.96 and an MCC of 0.61, suggesting it as a robust LBCE predictor for these viruses. Other PITL(ESM2)-based models with high performance gains include the predictors derived for two bacteria - *E. coli* (*AUC* = 0.91; *MCC* = 0.44) and *C. trachomatis* (*AUC* = 0.83; *MCC* = 0.57) - and one eukaryotic species - *P. falciparum* (*AUC* = 0.79; *MCC* = 0.41), demonstrating the broad applicability of our strategy across phylogenetically diverse pathogens. These models, as well as the others explicitly labelled in Figure 2(C), were shown to provide LBCE predictions with considerably better results for their target taxa than all state-of-the-art predictors against which they were compared.

As expected, a few cases for which we would not recommend the PITL-based models derived in this work over existing approaches were also observed. Only a single dataset, *M. tuberculosis*, resulted in the PITL(ESM2)-based model showing lower AUC against all external baselines. This appears to represent a particularly challenging dataset, with all methods tested exhibiting poor AUC (0.52 for PITL(ESM2), 0.54 for Epitope1D, 0.55 for EpitopeVec, 0.59 for Epidope, and 0.61 for BepiPred 3). Other datasets for which the PITL(ESM2)-based models did not perform well were *B. pertussis* (AUC = 0.53) and *S. mansoni* (*AUC* = 0.54). For the two other datasets explicitly labelled in the negative gains part of Figure 2(C), the comparatively low performance against BepiPred 3 is a result more of that method’s excellent performance for those datasets, rather than of poor performance of the PITL(ESM2)-based model (see Supplementary Table 5).

## 3 Conclusions

We show that phylogeny-informed transfer learning substantially improves protein language model–based prediction of linear B-cell epitopes. To this end, we implemented a general modelling framework to instantiate taxon-specific LBCE predictors using phylogeny-informed transfer learning to fine-tune a protein language model, coupled with a model fitting approach to optimise the predictive model to any target pathogen or taxon of interest. While fine-tuning of protein language models is known to improve performance in a variety of downstream tasks (Schmirler et al., 2024), a systematic statistical evaluation of the specific contribution of phylogenetic information in this setting has been lacking, despite recent work incorporating phylogeny into genomic language models (Tule et al., 2025) and across-species transfer learning (Ovaskainen et al., 2025). By contrasting the performance of PITL-based models with models developed without the embedder refinement step, and without the consideration of evolutionary relationships at the PLM fine-tuning stage, we show that the observed performance gains are primarily attributable to the phylogeny-aware data selection for the refinement of the ESM protein language model. We also established that PITL-based LBCE predictors provide substantial expected performance gains in AUC (between +0.09 and +0.123) over both generalist LBCE predictors and a recent taxon-specific predictor.

In its current version, the proposed PITL framework can be used to generate bespoke models for the prediction of LBCEs for a broad range of pathogens, including viruses, bacteria, and eukaryotic taxa. This approach allows for a reasonably straightforward development of models optimised for both well-studied pathogens and, more critically, for emerging, reemerging or neglected infectious diseases, through the selection of the most appropriate lower-level taxon from a data availability perspective. A notable limitation is the relative scarcity of well-curated LBCE datasets for fungal pathogens, which currently constrains applicability in this group.

Although we used the epitope prediction problem to illustrate the applicability of PITL to refine predictive pipelines in computational biology by leveraging evolutionary relationships, the framework is not restricted to this application. In a broader methodological context, adapting the framework presented here to other supervised learning tasks featuring cross-species biological data is a straightforward task, and the results presented in this work suggest that evolutionary structure can be explicitly incorporated into representation learning with PLMs, leading to significantly improved predictive models tailored to specific taxa. Because phylogeny is one instance of hierarchically structured information, we expect that a similar strategy may also improve modelling tasks in other domains where hierarchical structure governs data relationships.

## 4 Methods

### 4.1 Modelling

The framework for instantiating taxon-specific predictors consists of three main components (see Figure 1). Given a pair of datasets for matched {higher, lower} level taxa, the model development tasks are as follows.

The first step is *Embedder development*, consisting of the fine tuning of a general-purpose PLM to the task of interest, which in this case consists of LBCE prediction (see Supplementary File 1, section “Embedder Development Details”). In this task, the PLM is re-trained using entries from the higher-level dataset, excluding the target taxon entries to prevent overestimation of model performance caused by the usage of sequences of the target taxon during the training step. This results in a fine-tuned PLM for that particular phylogenetic group. Through this step, the structure learned as part of the ESM model’s primary development (Hayes et al., 2024) is augmented with knowledge about the representation of LBCEs from phylogenetically-related pathogens. We used the 650M parameter versions of ESM-1b and ESM2 as base models; however, the modular nature of the PITL framework makes it simple to use different embedder model sizes or different types of PLM.

The second step is *Feature calculation*, in which the fine-tuned embedder is deployed to extract features for the sequences from the target (lower-level) taxon. In the feature generation process, full protein sequences are fed to the refined PLM instead of just the labelled peptides, which enables the embedder to capture richer, non-local contextual information. This is expected to generate enhanced feature representation for each residue position. The output of this feature calculation model is then reduced so that only the labelled peptide regions are selected for later classifier training and performance assessment.

The third step is *Predictive model training and optimisation*, in which the features generated in the previous step are used for model fitting and hyperparameter tuning, resulting in a bespoke LBCE predictor for the target taxon. In this work, a simple Random Forest classifier (Breiman, 2001) was used to develop the final predictor, following the modelling approach used in previous works (Ashford et al., 2021; Campelo et al., 2024).

Supplementary File 1 provides further details on the network architecture of the ESM models and the details of the model and hyperparameter tuning of the classification models.

### 4.2 Performance assessment and comparison

All performance values presented in this work correspond to results estimated using *out-of-sample* predictions (i.e., data not seen at any point during embedder fine-tuning or model development). For the PITL-based models and the internal baselines, this represents an independently extracted test set for each target taxon (referred to in the text as “*Internal holdout*”). Supplementary File 1, section “Training and Validation Procedures”, provides details on the data splitting used and other aspects of model development and assessment.

Since peptides in the *Internal holdout* sets are not necessarily outside the training data of the different external baselines used, we assessed these tools primarily using observations of each target taxon not found in these methods’ published training datasets (referred to as “*No-leakage holdout*”). This approach is more likely to provide unbiased estimates of their generalization performance for each target taxa used in our experiments. Additionally, we also calculated the performance of the external baseline methods on the *Internal holdout* sets. These are reported alongside the main results in section 2.

To account for the effects of different methods, ESM versions, and datasets, we compared the performance of PITL-based models against the internal baselines using a factorial analysis of variance (ANOVA) design, with Method and Embedder as experimental factors, and Dataset as a blocking factor (Montgomery, 2017). Post-hoc pairwise tests for the differences of mean performance of different methods (accounting for Embedder and Dataset effects) were conducted using Dunnet’s all-vs-one contrasts (Montgomery, 2017), with PITL as the reference level for comparison.

The predictive performance of all models was quantified using the Area Under the ROC curve (AUC) as the main performance indicator, to ensure comparability with existing literature. Besides AUC, we also report eight additional performance measures in Supplementary Tables 3–5.

## 7 Data availability

The source code, data and analysis scripts used for this manuscript are deposited in Zenodo at https://doi.org/10.5281/zenodo.15743750. The development version of the code can be found at https://github.com/fcampelo/epitopetransfer.

## 8 Author contributions statement (CRediT)

L.P.L.: Software, Validation, Methodology, Writing (original draft); T.E.C.: Supervision, Resources, Writing (review & editing); F.P.L.: Conceptualization, Writing (Review & Editing); F.C.: Conceptualization, Methodology, Data curation, Visualization, Supervision, Writing (original draft), Writing (Review & Editing). All authors read and approved the final manuscript.

