## Supplementary files, figures and tables for "Phylogeny-informed transfer learning with protein language models for epitope prediction": Supplementary figure captions.docx

**Supplementary Figure 1: Fine-tuning data for phylogeny-informed (PITL) and phylogeny-agnostic (PATL) transfer learning**

Set of groups used for transfer learning. PITL uses a phylogenetic filter to select the data used for PML fine tuning. PATL, which uses biological groups unrelated to the target taxon, is used to test hypotheses about the effectiveness of using a phylogeny-informed strategy for transfer learning. If the PATL baseline models had produced equivalent or favourable results in relation to PITL, this would have indicated that enhancements observed in epitope prediction tasks could be exclusively attributed to the effect of fine tuning the representation to the predictive task, without any need to consider evolutionary relationships. Instead, the favourable results observed for PITL-derived models provide compelling evidence for the presence of a positive effect of using phylogeny as an explicit criterion when selecting data for fine-tuning the PML embedder.

**Supplementary Figure 2: Data consolidation step.**

This is an illustrative example of how the data consolidation step treated overlapping (and sometimes contradictorily labelled) peptides. Entries were aggregated in a residue-by-residue manner. Labels were attributed based on the mode (most common value), with ties being removed. “Label summary” summarises the total number of positive/negative labels on each region of the protein for this mock example, which was then used to determine the final labels of the consolidated entries.
