## Supplementary files, figures and tables for "Phylogeny-informed transfer learning with protein language models for epitope prediction": Supplementary File 1.docx

**Phylogeny-informed transfer learning with protein language models
for epitope prediction**

### Data

#### Data extraction and preparation

Data extraction, filtering, and consolidation were performed using the *epitopes* R package [1], according to the procedures outlined below.

Epitope data was retrieved from the complete XML export of the Immune Epitopes Database, IEDB [2]. All entries classified as related to linear B-cell epitopes (LBCEs) from organisms within the taxa Viruses (NCBI:txid10239), Bacteria (NCBI:txid2), and Eukaryota (NCBI:txi2759) were extracted from the IEDB export. The proteins associated with each entry were retrieved from either the NCBI protein database [3] or UniprotKB [4], based on the protein IDs provided in the metadata of each IEDB record.

To improve data quality, a quality control step was performed according to the following steps:

- Each entry retrieved from the IEDB was labelled as either a positive (i.e., corresponding to an epitope-containing region) or negative (a non-epitope containing region) peptide, based on a simple majority of the results of the assays reported for that entry.
- Positive-labelled peptides of length shorter than 5 or greater than 30 residues were removed to prevent very short entries or long “Epitope-containing regions” from adding excessive noise to the training data. This range was determined as a broader interval than the usual range of 15–25 residues for LBCE lengths [5,6] to strike a balance between data quality and excessive data removal. Negative-labelled entries were not filtered for size, since it is possible to have arbitrarily short or long sequences containing no epitopes.
- All entries were checked for positional consistency in their source proteins. Cases for which the expected coordinates on the protein did not perfectly match the peptide were discarded.

After quality control, the remaining labelled peptides were projected onto their source protein sequences and checked for overlaps. Where overlaps occurred, residues were labelled as the mode of the labels attributed to that position, with ties being assigned an unlabelled status. Finally, the entries were consolidated such that adjacent labelled regions with the same label were merged into a single labelled peptide. Supplementary Tables 1-2 summarise the data volume under each dataset. Supplementary Figure 2 illustrates the data consolidation step.

Entries were clustered based on source protein dissimilarity, calculated using DIAMOND [7], using agglomerative clustering with single linkage. To account for possible local (peptide-level) similarities, any alignments with coverage greater than 8 were considered, and a 30% similarity threshold was applied to define the groupings. Clusters were treated as the basic splitting unit when isolating the final test sets (referred to as *Internal holdout* in the text), as well as for cross validation and hyperparameter tuning, to minimize data leakage due to similarity/homology.

To investigate the transfer learning approach proposed in this work, we instantiated nineteen pairs of datasets (higher- and lower-level) for a diverse range of pathogens, including bacterial, viral and eukaryotic pathogens. Supplementary Tables 1-2 provide the details of the datasets used, including the number of positively- and negatively-labelled peptides at each level, the taxonomic levels used for each case, and the full list of NCBI taxonomy IDs encompassed by each dataset.

The criteria for selecting the lower taxonomic levels were potential clinical interest and diversity of taxonomic levels. Pathogens of clinical importance that researchers could be interested in exploring were initially selected at the species level. If that species had sufficient data for model training (defined as more than 10 labelled peptides for each class, based on the preliminary findings reported in [9]) it was set as the lower-level taxon. Otherwise, the taxonomic level was increased until a sufficient data volume was achieved.

Once the lower-level taxon was determined, its higher-level counterpart was defined as the lowest level that contained a substantial number of labelled peptides (not including the lower-level entries), sufficient for the ESM model fine tuning. This was defined as at least 200 labelled peptides (with one exception made for the Orthopoxvirus-Bamfordvirae pair, in which the labelled data volume at the kingdom level is still slightly short of 200 labelled peptides).

### Embedder Development Details

The embedder development process has two main components: ESM fine-tuning and feature calculation. A key aspect to fine-tuning step is the use of a sliding window to compute amino acid level data.

#### Sliding window

A sliding window was implemented to deal with the limitation of sequence lengths emerging from the maximum capacity permitted by the ESM model (1024 residues). For protein sequences longer than that, a sliding window was implemented with length 1024 and a step size 512. The window was truncated if it exceeded the sequence length The ESM embeddings were calculated for each these sub-sequences and later aggregated using per-residue averaging.

#### ESM fine-tuning

We proceed to fine-tune the ESM model by integrating a classification head into the original architecture, as detailed in subsection “ESM Model Architecture”. This fine-tuning involves a retraining of all layers of the model. Thus, the initial model trained for general protein sequence analysis is now tailored to epitope prediction task. To this procedure, the dataset comprises sequences from higher-level taxa, as outlined in section “Data extraction and preparation” of the main text and detailed in Supplementary Tables 1-2. The fine-tuning process leverages the ESM models’ capabilities to learn from our specialized dataset, focusing on epitopes and non-epitopes present in these sequences. In the fine-tuning phase, the ESM model processes each training sample, extracted through the sliding window technique. The fine-tuning is executed over three epochs, and the model learns to identify and differentiate between epitopes and non-epitopes in the context of the higher-level taxa sequences. The low number of epochs was used to reduce the chances of model overfitting, given the relatively modest data volumes available for fine-tuning (when compared to the number of model parameters).

The result of this procedure is a fine-tuned model specifically capable of identifying epitopes in protein sequences derived from higher-level taxa. This model is now able to generate enriched features to epitope prediction classification tasks.

#### Feature Calculation

In this step, we employ the fine-tuned ESM model, which has been pre-trained with data from higher taxonomic levels, to process the amino sequences and generate enriched features to epitope prediction task at lower taxonomic levels. The complete protein sequence is fed into the optimized ESM model, enabling it to generate a representation that accounts for both local and non-local interactions. This 1280-dimensional feature representation encodes the properties and contextual information that the model has learned. After the feature calculation step, the positions corresponding to labelled peptide regions are extracted for training.

#### ESM Model Architecture

To enable the fine tuning of ESM embedders in this work, we added a “Classification Head” layer to the original model architecture, which is later removed to enable the extraction of features from the taxon-optimised model. The architecture can be summarized by its key components, which include:

**Embeddings:**

- *Word Embeddings:* Convert each residue in a sequence to a 1280-dimensional vector. The word embeddings are **trainable**, meaning their values are adjusted during model training to improve task-specific performance.
- *Position Embeddings:* Provide positional context to each residue in sequences, up to 1024 positions long. Unlike the word embeddings, the position embeddings are **fixed** and are not updated during training, ensuring that the positional information remains consistent.

**Encoder** comprising 33 layers, each featuring:

- *Self-Attention Mechanism:* Allows each position to interact with every other position in the sequence.
- *Feed-Forward Network:* Enhances the model output from the self-attention mechanism by applying transformations.
- *Layer Normalization:* Applied for training stability.

**Contact Prediction Head:**

- A specialized component for predicting protein structure contacts. This layer involves determining which pairs of amino acids within a sequence are in close proximity to each other in the folded structure, referred to as contacts.

The ESM model, tailored for protein sequence analysis, comprises a 33-layer encoder, where each position in the protein sequence is represented by a 1280-dimensional vector. While the model supports an embedding size of 1026, the practical window for sequence analysis utilizes 1024 positions. This is due to the allocation of one special token at the beginning and another at the end of the sequence.

### Training and Validation Procedures

*Data splitting and cross validation***:** For each taxon, the data is initially split into five subsets, using the similarity-based groupings described in Section ”Data” to prevent data leakage across folds. One such subset is set aside for final performance assessment, and the remaining four ones are used for model development and validation.

A nested cross-validation is performed using these four folds: three folds are used for training (3-fold CV) and one for validation, such that the models after the hyperparameter tuning step are assessed on unseen data.

*Hyperparameter Search***:** Random Forest hyperparameters are optimized using Bayesian optimization as implemented in the *Optuna* package [8]:

- estimators (range: 100 to 500)
- maxdepth (choices: None, 10, 20, 30)
- minsamples split (range: 2 to 10)
- minsamples leaf (range: 1 to 10)
- maxfeatures (choices: sqrt, log2, None)
- bootstrap (choices: True, False),
- criterion (choices: gini, entropy)

The best hyperparameters are defined during this process. After this step, the model is retrained using all four folds and the best hyperparameters found, and the optimal threshold (the one that results in the highest MCC) is determined. Finally, the model's performance is evaluated on the test set.

#### Special Cases

Due to the scarcity of samples for *Orthopoxvirus*, *Corynebacterium*, and *Measles morbilivirus*, we followed a standard train-test procedure for these cases instead of cross-validation. For each taxon, the data was split in two folds. One was set aside for testing, and the remaining fold was used for training. In these scenarios, no hyperparameter optimization was conducted; instead, the Random Forest classifier from scikit-learn was implemented using its default hyperparameter settings, as summarized below:

- *n estimators* = 100
- *criterion* = gini
- *max depth* = None
- *max features* = sqrt
- *min samples split* = 2
- *min samples leaf* = 1
- *bootstrap* = True

The classification threshold in these two cases was kept at 0*.*5.

### End-to-End Neural Network and other classifier alternatives

The current hybrid approach combines a fine-tuned ESM embedder with a Random Forest classifier. The framework presented in this paper is, however, model-agnostic, and could easily be adapted to use different classifiers such as XGBoost, Bayesian classifiers, Logistic regression, or even a feed-forward neural network. In this last case, the Random Forest would be replaced by a feed-forward neural network (FFNN) classification head attached directly to the ESM encoder, enabling end-to-end training via backpropagation.

A key challenge in using a full neural network-based approach lies in determining the extent of neural network optimization required to maximize performance while preventing overfitting, as the optimal stopping point may vary considerably across different taxa. In contrast, the Random Forest model employed in our experiments is a much simpler model, and less susceptible to overfitting. Users of the proposed method can, however, adapt it using different classifiers or feature embedders, depending on their specific requirements and preferences.

### Computational resources

Fine-tuning the ESM models in our experiments required 4 GB RAM, 30 GB GPU RAM, and 44 GB disk space. For prediction generation with Random Forest, we used a standard machine with 36 GB RAM, 30 GB disk space, and 8 CPUs.

### Performance Indicators

A range of performance indicators was used to estimate model performance. These metrics not only allow for a comparison with existing and future studies but also provide a detailed insight into the predictive accuracy of the models under consideration. The definitions of these indicators involve key terms such as TP (True Positives), TN (True Negatives), FP (False Positives), and FN (False Negatives), which are fundamental to the formulae that follow.

- **Positive Predictive Value (PPV)**: probability that a residue predicted as positive is indeed part of an epitope sequence. It serves as an indicator of the reliability of a model’s positive predictions. The Positive Predictive Value (PPV) is commonly recognized in the field as Precision.

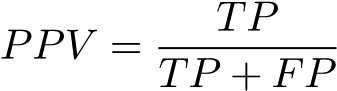

- **Negative Predictive Value (NPV)**: probability that a residue predicted as negative is indeed not part of an epitope sequence. It serves as an indicator of the reliability of a model’s negative predictions.

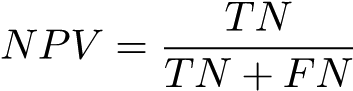

- **Sensitivity (SENS)**: Sensitivity, also called Recall or True Positive Rate (TPR), measures the model’s ability to correctly identify residues in known epitope sequence.

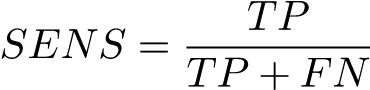

- **Specificity (SPEC)**: Specificity, or the True Negative Rate (TNR), measures the model’s ability to correctly identify residues in known non-epitope sequence.

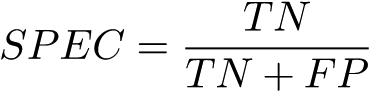

- **Accuracy (ACC)**: This overall metric represents the rate of correct classifications made by the model. While providing a general indication of performance, it should be noted that accuracy can be influenced by class imbalance and does not always reflect the differential costs associated with false positives and false negatives, especially in epitope prediction contexts.

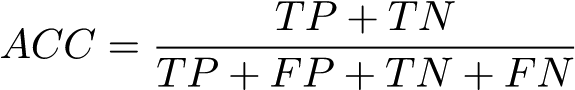

- **AUC (Area Under the ROC Curve):** This indicator provides an assessment of a classifier’s performance across different thresholds.

$$AUC=\int_{0}^{1} C\left( x \right)dx$$

Where x *C(x)* denotes the curve of the false positive rate (*x*, defined as 1 - Specificity) versus Sensitivity, derived by varying the classification threshold from zero and one.

- **F_1_ Score:** This is a metric that balances Precision and Recall, making it useful for evaluating binary classification models, especially when dealing with imbalanced datasets. It combines Precision (the ability to correctly identify positive samples) and Recall (the ability to capture all positive samples) into a single score.

$$F_{1}= \frac{2\times PPV\times SENS}{PPV+SENS}$$

- **MCC (Matthews Correlation Coefficient):** The Matthews Correlation Coefficient (MCC) is a metric that considers True Positives (TP), True Negatives (TN), False Positives (FP), and False Negatives (FN) to assess the performance of binary classification models. It produces values between -1 and +1, where +1 indicates perfect prediction, 0 corresponds to random prediction, and -1 reflects total disagreement between predictions and actual outcomes. MCC is particularly useful when dealing with imbalanced datasets.

$$MCC= \frac{TP\times TN-FP\times FN}{\sqrt{(TP+FP)(TP+FN)(TN+FP)(TN+FN)}}$$
