## Supplementary figures and images for "Phylogeny-informed transfer learning with protein language models for epitope prediction"

### Supplementary Figure 1.png

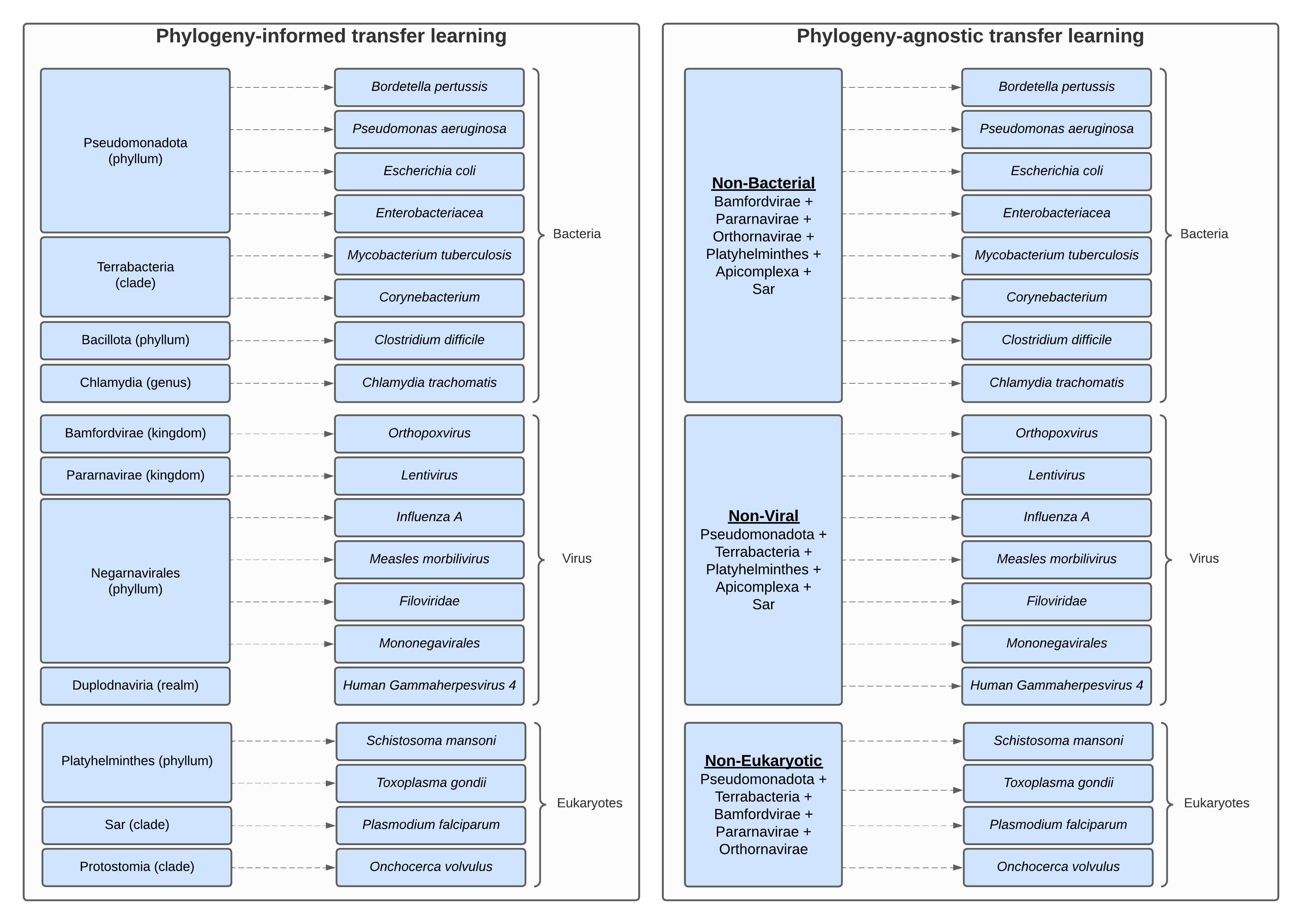

### Supplementary Figure 2.jpg

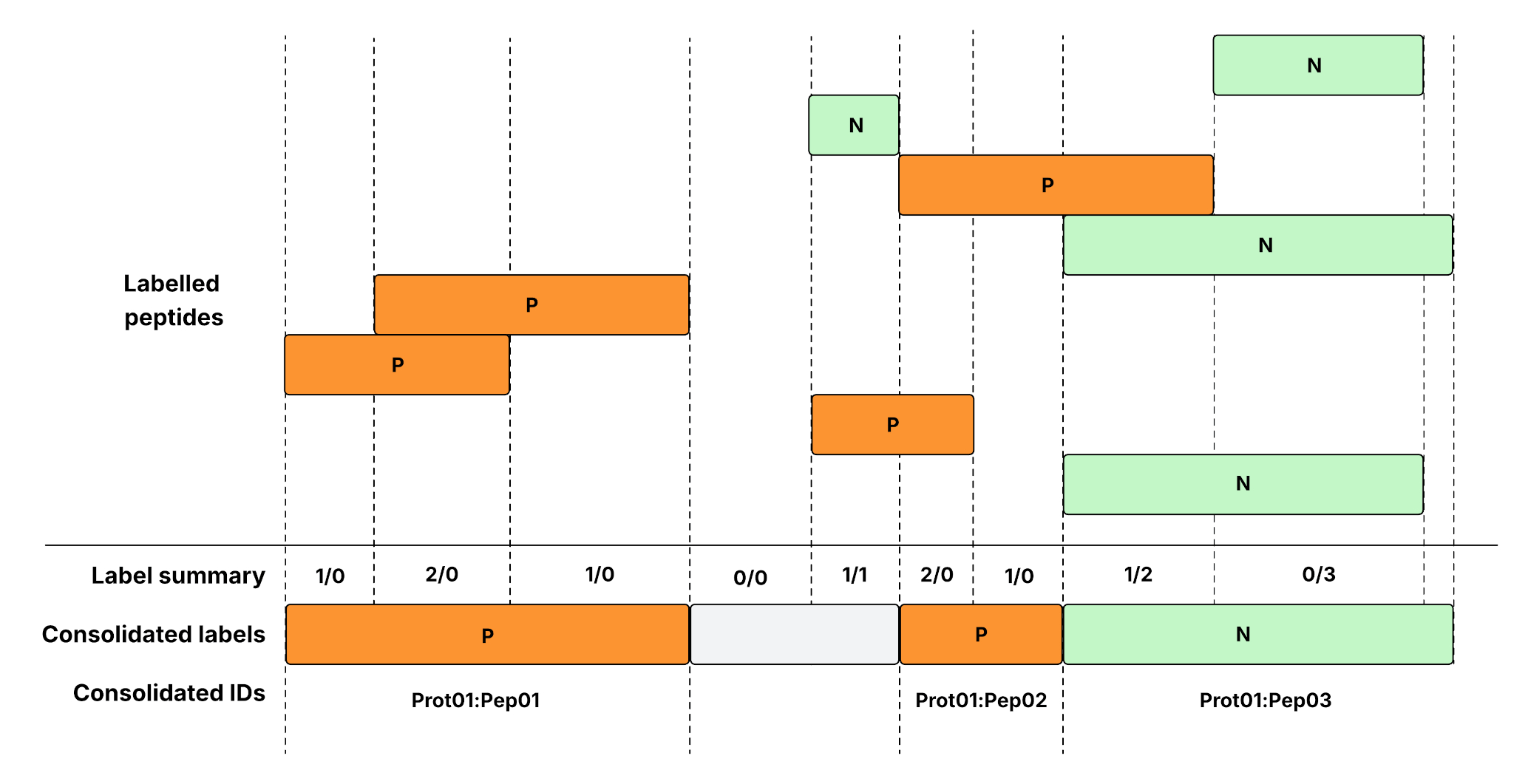
